# Development of Polarity-Reversed Endometrial Epithelial Organoids

**DOI:** 10.1101/2023.08.18.553918

**Authors:** Vakil Ahmad, Sai Goutham Reddy Yeddula, Bhanu P. Telugu, Thomas E. Spencer, Andrew M. Kelleher

**Affiliations:** Division of Animal Sciences, University of Missouri, Columbia, MO, 65211; Department of Obstetrics, Gynecology, and Women’s Health, University of Missouri, Columbia, MO, 65211

**Keywords:** Uterus, Epithelium, Organoid, Polarity

## Abstract

The uterine epithelium is composed of a single layer of hormone responsive polarized epithelial cells that line the lumen and form tubular glands. Endometrial epithelial organoids (EEO) can be generated from uterine epithelia and recapitulate cell composition and hormone responses *in vitro*. As such, the development of EEO represents a major advance for facilitating mechanistic studies *in vitro*. However, a major limitation for the use of EEO cultured in basement membrane extract and other hydrogels is the inner location of apical membrane, thereby hindering direct access to the apical surface of the epithelium to study interactions with the embryo or infectious agents such as viruses and bacteria. Here, a straightforward strategy was developed that successfully reverses the polarity of EEO. The result is an apical-out organoid that preserves a distinct apical-basolateral orientation and remains responsive to ovarian steroid hormones. Our investigations highlight the utility of polarity-reversed EEO to study interactions with *E. coli* and blastocysts. This method of generating apical-out EEO lays the foundation for developing new *in vitro* functional assays, particularly regarding epithelial interactions with embryos during pregnancy or other luminal constituents in a pathological or diseased state.

## Introduction

Understanding uterine epithelial biology is essential for improving women’s reproductive health, as defects in epithelial function contribute to various uterine pathologies and infertility (Gruber and Mechsner, 2021, Wang et al., 2020, Burney and Giudice, 2012, Strauss and Barbieri, 2019). The uterine epithelium consists of a single layer of pseudostratified luminal epithelium and simple columnar glandular epithelium. The uterine epithelium has a critical role in maintaining the contents of the uterine lumen, facilitating blastocyst adhesion and attachment, and protecting the uterus from ascending infections (Kelleher et al., 2019, Mitchell et al., 2015, Moreno et al., 2016, Benner et al., 2018, Aplin and Kimber, 2004). However, our understanding of the cellular and molecular mechanisms underlying endometrial epithelial physiology and function is limited due to the technical constraints of *in vitro* models and the ethical limitations of *in vivo* investigations particularly during pregnancy.

Organoids have recently emerged as an indispensable model for studying epithelial biology, pathophysiology, and embryo-maternal interactions (Murphy et al., 2022, Fitzgerald et al., 2021, Fitzgerald et al., 2019, Boretto et al., 2019, Turco et al., 2017, Boretto et al., 2017). To establish endometrial epithelial organoids (EEO), epithelial cells from the endometrium are isolated through enzymatic digestion and mechanical disruption. These cells are then embedded within a basement membrane extract (BME) and form three-dimensional structures. Growth media supplemented with a cocktail of growth factors promote cell proliferation, differentiation, and self-organization into mature organoids (Turco et al., 2017, Boretto et al., 2017). Importantly, EEO can be propagated while retaining key features of the endometrial epithelium, including tissue organization, cellular composition, gene expression signatures, steroid hormone response, and protein secretion profiles (Fitzgerald et al., 2019, Turco et al., 2017, Boretto et al., 2017, Fitzgerald et al., 2023). Thus, EEO have the capacity to serve as a powerful in vitro model with significant potential for addressing key questions in reproductive biology related to epithelial function, regeneration, and development.

A defining feature of BME-embedded three-dimensional organoid structures is a polarized epithelium with a central lumen. The apical-in/basal-out polarity of EEO presents a challenge to study epithelial interactions with embryos or infection by microbes or viruses that reside in the uterine lumen (Boretto et al., 2019, Boretto et al., 2017, Co et al., 2019, Simintiras et al., 2021, Kakni et al., 2023, Wijesekara et al., 2023). To overcome this limitation, a suspension culture method to reverse EEO polarity was developed based on methods reported for human enteroid and airway organoids (Co et al., 2019, Co et al., 2021, Kakni et al., 2023, Wijesekara et al., 2023). Our studies demonstrate that this culture method alters the polarity of EEO without compromising their physiological hormone response and viability. The polarity reversed EEO exhibit histological and physiological characteristics resembling uterine epithelium *in vivo*, respond to hormones, and undergo secretory cell transformation. Co-culture experiments further reveal the preferential interaction of microbes with polarity-reversed organoids, providing a more representative *in vivo* condition. Likewise, co-culture with blastocysts demonstrates the hormonal impact on EEO-embryo interactions. The apical-out EEO can be utilized for a broad range of applications beyond the scope of this initial characterization, including studies on epithelial-microbial interactions, drug screening, and investigations into uterine epithelial-embryo interactions during implantation.

## Results

### Development of endometrial epithelial organoids

The current EEO model poses challenges in studying experimental interactions between the epithelial surface and luminal contents, such as during embryo-uterine interactions and microbial infections. Building on the concept of polarity reversal observed in organoids derived from the gastrointestinal tract (Co et al., 2019, Co et al., 2021, Kakni et al., 2023, Wijesekara et al., 2023), we determined if the polarity of EEO could be reversed by chemically releasing intact organoids from the basement membrane extract (BME) and subsequently cultivating them in suspension culture. These studies employed EEO from the endometrium of healthy women that were cultured in BME under WNT activating growth conditions as described previously (1–3) (Fig. 1A). The human EEO formed spherical structures with the basolateral epithelial surface facing outward in contact with the BME, while the apical surface was oriented towards the inside of the organoid (Fig. 1B). To maintain the integrity of EEO during removal from BME, cell recovery solution was used to depolymerize the BME and release organoids without disrupting the organoid three-dimensional structure. The EEO were then transferred to suspension culture in growth media using ultra low-attachment plates (Fig. 1A). Visible changes in the gross morphology of EEO was observed in suspension culture, but both BME-embedded and suspension cultured EEO maintained a single-layered epithelium with a central lumen (Fig. 1B; c, d). Immunofluorescence imaging of apical surface proteins zonula occludens 1 (ZO1) and F-actin revealed that EEO in suspension culture exhibited reversal of apical-basal polarity after several days in culture, resulting in an outward-facing apical surface (referred to as apical-out EEO or AO). In contrast, BME-embedded organoids (referred to as apical-in EEO or AI) displayed the expected apical surface facing the central lumen (Fig. 1B; e-f and Fig S1A). Scanning electron microscopy (SEM) further confirmed the polarity reversal with the outward apical surface covered with microvilli in suspension cultured organoids, while the outward surface of AI organoids was smooth (Fig. 1B; g, h). Importantly, the viability and growth of EEO in suspension culture was not altered with similar numbers of proliferating cells based on Ki67 immunostaining compared to BME-embedded EEO (Fig S1B).

**Figure 1:**
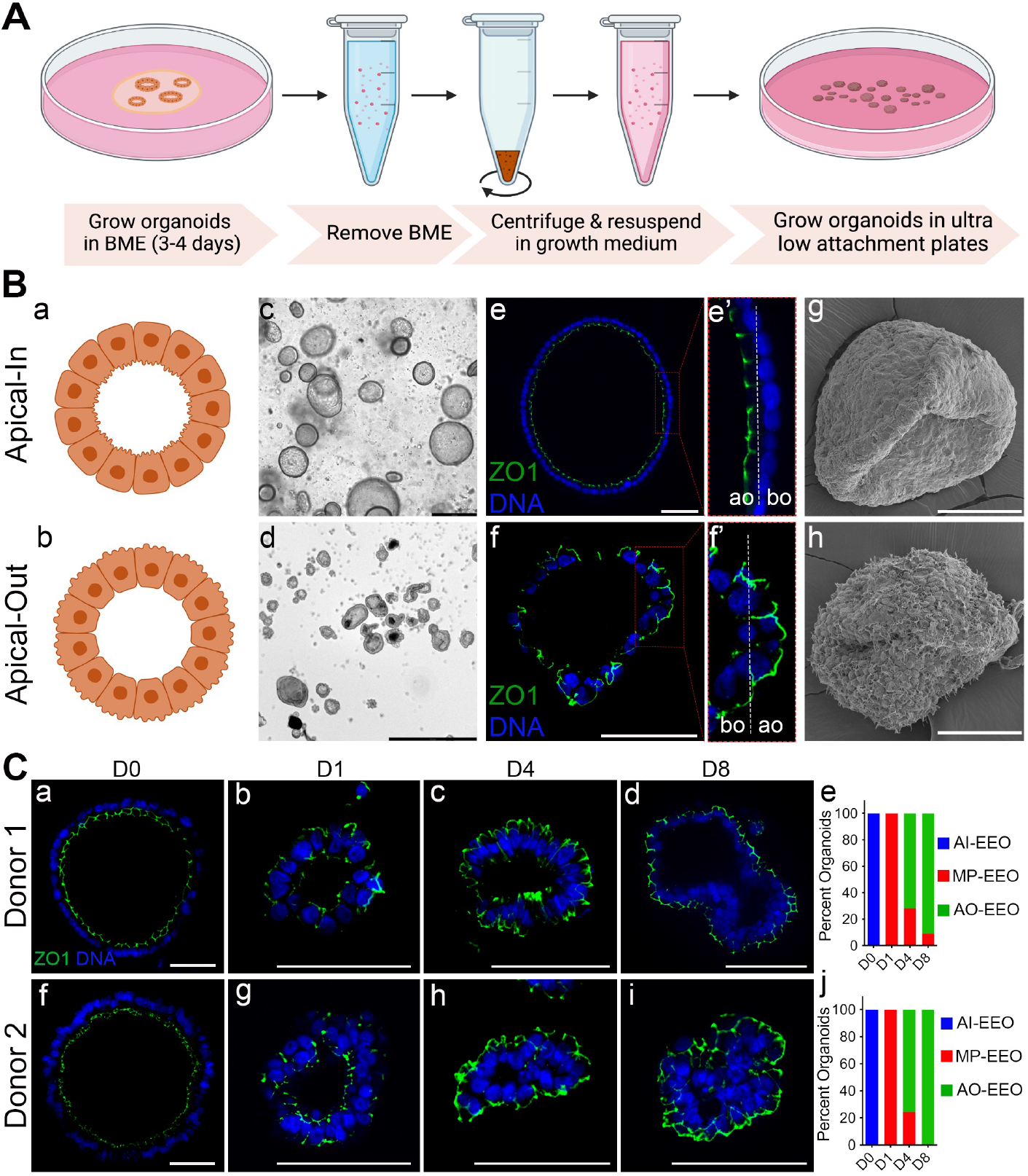
Generation and characterization of apical-out organoids. (**A**) Approach used to generate apical-out endometrial epithelial organoids (AO-EEO or AO); (**B**) Characterization of AO. (**a-b**) illustration of apical-in organoids and AO EEO; (**c-d**) bright field images of organoids grown in Cultrex BME or in suspension culture for eight days (scale bars, 100 µm); (**e-f**) Immunofluorescence of the apical surface marker ZO1 in AO or AI (scale bars, 20 µm); (**e’-f’**) Areas outline d in boxes in **e** and **f** magnified to demonstrate the apical localization of ZO1; (**g-h**) SEM analysis of AI or AO (scale bars, 50 µm). (**C**) Time-course analysis of EEO polarity reversal from day 0 to day 8 in suspension culture; (a-d; f-i) The apical surface is detected by ZO1 immunofluorescence for Donor 1 and Donor 2 (scale bars, 50 µm); (**e and j**) Quantification of the percentage of AI and AO organoids in the time course experiment for Donor 1 (e) and Donor 2 (j) (scale bars, 50 µm).

To understand the kinetics of EEO polarity reversal, immunofluorescent microscopy was used to quantify the percentage of EEO exhibiting apical-in, apical-out, or mixed (partial apical-in and apical-out) polarity in a time course experiment. EEO obtained from two independent donors were released from BME, suspension cultured, and evaluated over 8 days. ZO1 localization revealed the reversal of apical-basal polarity commenced within 24 hours of the initiation of suspension culture, with all EEO displaying mixed polarity after on day 1 (Fig. 1C; a, b, f, g, e, j). On day 4 of suspension culture, the majority (>70%) of EEO exhibited apical-out polarity (Fig. 1C; c, h, e, j). On day 8, over 90% of EEO from both donors demonstrated AO polarity (Fig. 1C; d, i, e, j). SEM analysis found that microvilli emerged on the outer surface of EEO cultured in suspension by day 1 with increased abundance observed on days 4 and 8 (Fig. S1C). Of note, this developed polarity reversal method was also successful with mouse EEO, resulting in complete polarity reversal in suspension culture (Fig. S1D). These results indicate that EEO polarity reversal can be reliably achieved during one week by removing BME and performing suspension culture.

### Apical-out EEO exhibit physiological hormone responsiveness

EEO mimic physiological responses to progesterone including down-regulation of PGR (progesterone receptor) expression and up-regulation of progesterone-responsive genes (Fitzgerald et al., 2021, Fitzgerald et al., 2019, Turco et al., 2017, Fitzgerald et al., 2023). To assess the impact of polarity reversal on EEO response to steroid hormones, AI and AO EEO were treated with E2 followed by E2+MPA (Fig. 2A) as previously described (Fitzgerald et al., 2023, Fitzgerald et al., 2019). MPA is non-metabolizable PGR agonist. Both AI and AO responded to steroid hormone treatment, and no differences were observed in the number of PGR-positive and ESR1-positive cells between AI and AO organoids (Fig. 2B and Fig. S2A). Treatment with E2 alone increased the number of PGR positive cells, which was attenuated by MPA in both AO and AI EEO (Fig. 2B-E). Treatment with E2 also increased the number of ESR1 positive cells and increased the number of proliferative Ki67 positive cells in both AO and AI EEO compared to the control (Fig. S2A-B). The organoid response to steroid hormone treatment is consistent with previous reports and reflects the *in vivo* response to E2 and progesterone (Fitzgerald et al., 2023, Fitzgerald et al., 2019, Lessey et al., 1988).

**Figure 2.**
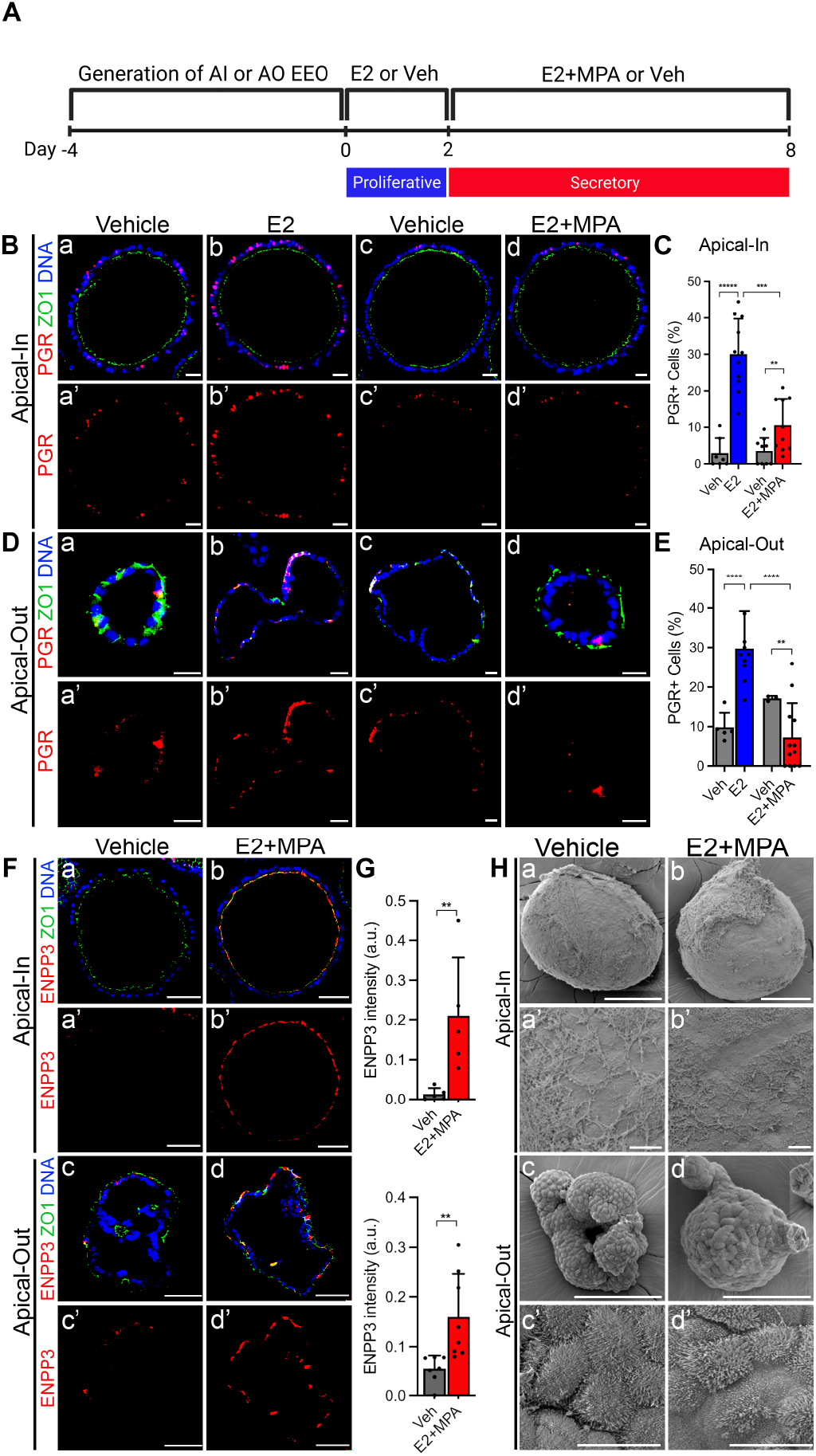
Steroid Hormones Elicit Similar Responses in Both Apical-In and Apical-Out Organoids. (**A**) Experimental design. (**B**) Representative images of immunofluorescence staining of PGR (red) and ZO1 (green) on AI or AO treated with vehicle or E2 for 2 days and/or E2+MPA for 6 days. (**C**) Quantification of PGR-positive cells. (**D**) Representative images of immunofluorescence staining of ENPP3 (red) and ZO1 (green) on paraffin sections of AI or AO organoids treated with vehicle or E2 for 2 days and then with E2+MPA for 6 days. (**E**) Quantification of ENPP3 (red) signal intensity. All scale bars in B, D & F are 20 µm. Plots in **C** and **E** represent mean ± SD. (**F**) Representative SEM images of AI or AO organoids treated with vehicle or E2+MPA showing the structural changes on organoids’ surface; scale bars, 100 µm (**a-d**). Magnified sections of ROI are shown; scale bar, 20 µm (**a’**-**d’**). \*\**p* < 0.01, \*\*\**p* < 0.001 and \*\*\*\**p* < 0.0001 (Student’s t test).

Ectonucleotide pyrophosphatase (ENPP3) is progesterone responsive gene in the human endometrium and EEO (Fitzgerald et al., 2023, Boggavarapu et al., 2016). ENPP3 was substantially increased in both AO and AI EEO treated with E2 and MPA (Fig. 2F-G). Immunoreactive ENPP3 was observed on the apical surface of the EEO, reflecting the *in vivo* secretion pattern of ENPP3 and the polarity reversal phenotype of organoids cultured in suspension (Fig. 2F). The apical surface of uterine epithelial cells undergoes significant changes in the appearance in response to steroid hormones and preparation for implantation (Murphy, 2004). SEM analysis of hormone-treated organoids revealed a decrease in the height and flattening of microvilli compared to the vehicle alone controls (Fig. 2H). Collectively, these results support the idea that AO EEO maintain their physiological responses to steroid hormones.

### Apical-out EEO for the study of cell interactions

EEOs offer significant potential for studying the interactions between the epithelium and components of the uterine lumen. In order to establish effective coculture systems for future mechanistic studies, interactions of AO EEO with a fluorescently labeled reporter strain of *E. coli* or mouse blastocysts were studied (Fig. 3A). An increased attachment of *E. coli*^*GFP*^ (ATCC #25922) was observed in AO organoids compared to AI organoids at 48 h post-infection (Fig. 3B, C and D). Extended culture for 96 h post-infection resulted in a steady increase in GFP intensity in AO organoids, indicating bacterial replication (Hirako et al., 2022). In contrast, the GFP intensity remained low and unchanged in AI organoids after 48 and 96 h post-infection (Fig. 3B, C and D).

**Figure 3.**
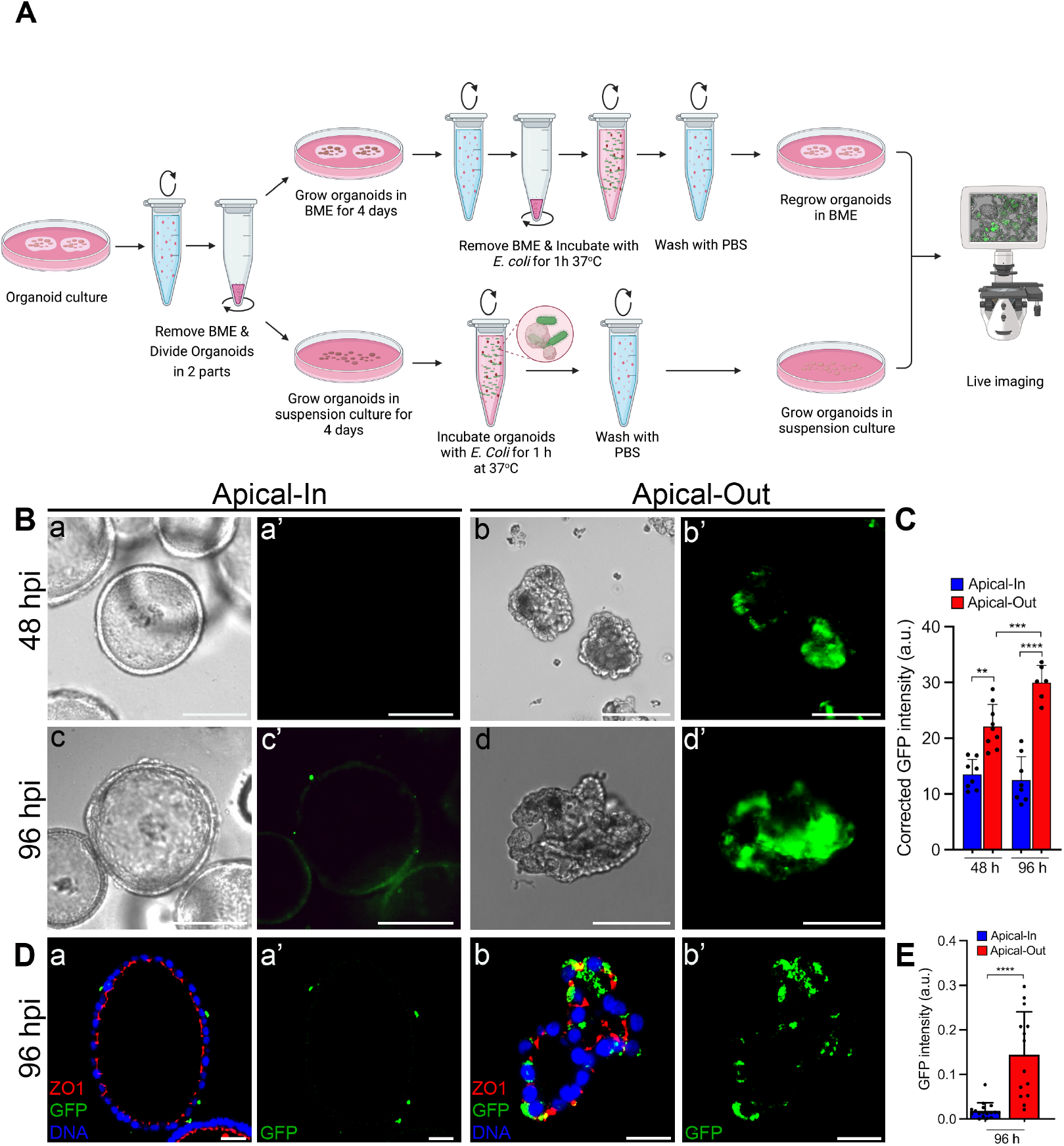
Apical-Out Organoids Exhibit Vulnerability to E. coli Infection. (**A**) Schematic of the experimental design to infect AI or AO with *E. coli*^*GFP*^. (**B**) Epifluorescence imaging of AI or AO organoids infected with GFP-labeled *E. coli*^*GFP*^ at 48 and 96 hours post infection (hpi); scale bars, 100 µm. (**C**) Quantification of GFP fluorescence intensity from (B). (**D**) Representative images of immunofluorescence staining of ZO1 (red) and GFP (green) infected with GFP-labeled *E. coli* at 96 hpi; scale bars, 20 µm. (**E**) Quantification of GFP fluorescence intensity from (D). Plots in **C** and **E** represent mean ± SD. \*\**p* < 0.01, \*\*\**p* < 0.001 and \*\*\*\**p* < 0.0001 (Student’s t test).

Finally, the co-culture of blastocysts with AO EEO demonstrated the potential for studying endometrial-embryo interactions without using BME (Fig. 4A). AO EEO were cultured in low attachment plates with implantation-competent mouse blastocysts expressing GFP (RRID: IMSR_JAX:004353) to evaluate the impact of steroid hormone treatment on EEO-embryo attachment. Blastocysts did not attach to vehicle treated AO EEO, whereas blastocysts did attach to AO EEO treated with a steroid hormone regimen mimicking the mid-secretory phase (Fitzgerald et al., 2023, Fitzgerald et al., 2019) (Fig. 4A and B). These findings confirm the ability of hormonally primed organoids to interact with and support the attachment of blastocysts (Rawlings et al., 2021), suggesting that this model could be an excellent tool for studying endometrial embryo interactions.

**Figure 4.**
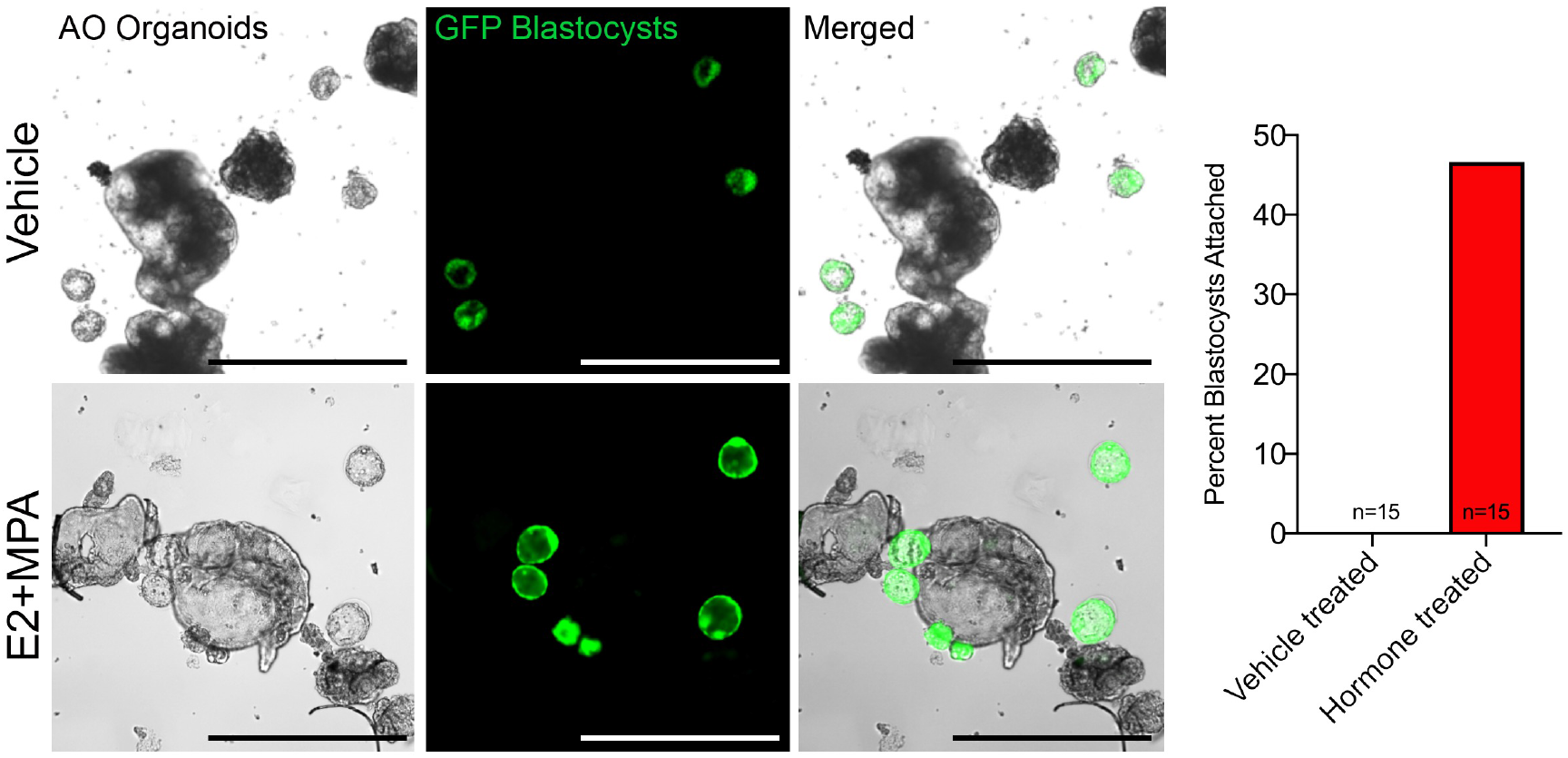
Blastocysts Attach to Hormone Treated Apical-out Organoids. (**A**) Bright field, GFP epifluorescence and merged images of AO treated with vehicle or E2 for 2 days and E2+MPA for 6 days, and then co-cultured with GFP labeled blastocysts for 24 h. 15-20 blastocysts were co-cultured with ∼50 vehicle treated or hormone-treated AO organoids from two independent donors. scale bars, 100 µm. (**B**) Quantification of blastocyst attachment in (A).

## Discussion

EEO are having a major impact on *in vitro* models of the uterus. These organoids have found broad application in studies pertaining to epithelial development, physiology, and disease due to their ability to mimic the properties of the uterine epithelium *in vivo*. Original reports and subsequent studies highlight EEO have a self-organized apicobasal structure, where the basal surface contacts the BME and the apical side is inside forming a closed lumen. Thus, access to the apical surface of basal-out EEO is restricted and microinjection techniques are required to deliver to or remove factors from the apical surface or lumen of the organoids (Co et al., 2019, Simintiras et al., 2021, Bartfeld and Clevers, 2015, Bartfeld et al., 2015). This technique can be technically challenging due to the requirement for organoid growth to large diameters and potential contamination of the apical organoid compartment. Moreover, normalizing the exposure of microinjected experimental agents into the organoid lumen is complex due to the small volumes and variability in organoid size.

Using techniques from other organoid systems (Co et al., 2019, Nash et al., 2021, Giobbe et al., 2021), polarity reversal was achieved with EEO from the human and mouse endometrium by simply removing BME and performing suspension culture for one week. The apical-out EEO model recapitulates critical functions of the endometrial epithelium *in utero*. Similar to basal-out EEO, apical-out EEO, despite the physical reorientation, retained their inherent functionality, including proliferation and steroid hormone response (Fitzgerald et al., 2023, Fitzgerald et al., 2019, Boretto et al., 2019, Turco et al., 2017, Boretto et al., 2017). Thus, polarity reversed EEO can withstand structural manipulation without compromising their biological properties, maintaining their usefulness in exploring endometrial physiology and pathophysiology (Fitzgerald et al., 2021, Boretto et al., 2019, Turco et al., 2017, Boretto et al., 2017). The ability to form these organoids with an apical-out orientation is expected to enhance their application in research that requires interaction between the luminal epithelium with embryos as well as microbes and viruses (Murphy et al., 2022, Fitzgerald et al., 2021, Simintiras et al., 2021).

To test the utility of the apical-out EEO model against the traditional apical-in model, a system was devised to investigate apical endometrial interactions with *E. coli* and mouse blastocysts. These serve as representative models for pathological and physiological interactions of the endometrial epithelium, respectively. The ability of the apical-out organoids to establish functional connections with these external entities suggests their potential use in investigating diverse interactions occurring within the uterine lumen. Specifically, the enhanced attachment and replication of *E. coli* on apical-out organoids compared to apical-in organoids present a useful model for studying ascending microbial infections. Such infections are key components of several reproductive tract diseases. Ascending *E. coli* infection from the vagina to the uterus is a significant cause of disease in both pregnant and non-pregnant women (Brunham et al., 2015). During pregnancy, *E. coli* infection can result in multiple negative consequences for the mother and child, such as neonatal or maternal sepsis, miscarriage, and pre-term birth (Cools, 2017). Further, mouse blastocysts would attach to apical-out organoids treated with steroid hormones. The direct study of human embryo implantation is challenging due to ethical concerns and technical barriers, and animal models do not perfectly mirror the human process. The successful co-culture and enhanced attachment of mouse blastocysts with apical-out organoids treated with steroid hormones marks an advancement in the potential to model *in vitro* endometrial-embryo interactions to study early pregnancy loss.

In summary, a validated method was developed here to generate an outward-facing endometrial epithelial surface in organoid culture. The demonstrated ability to modify the polarity of EEO without impairing their functionality, coupled with successful co-culture with microbial and embryonic entities, paves the way for investigating complex interactions at the endometrial epithelial surface at scale. These insights hold promise for elucidating physiological processes such as embryo implantation and pathogenic processes like infection. Future research can leverage this model to gain a deeper understanding of these interactions, potentially unveiling new therapeutic strategies for conditions related to the endometrial epithelium.

## Methods

### Endometrial Epithelial Organoid Establishment and Maintenance

Endometrial epithelial organoids (EEO) were isolated, cultured, and maintained as previously described (Fitzgerald et al., 2019, Turco et al., 2017, Fitzgerald et al., 2023). Briefly, primary human endometrial epithelial cells were isolated from endometrial biopsies of donors. Written informed consent was obtained from each donor and the study was approved by the University of Missouri Institutional Review Board (IRB Project Approval Number: 2011513). Those cells were washed and subsequently resuspended in Cultrex BME (Cat. No. 343300501, R&D Systems) and plated in 20-25 µl droplets of Cultrex. The droplets were solidified by incubation at 37°C for 15 minutes before being overlaid warm organoid expansion culture medium as reported previously (Fitzgerald et al., 2019). To passage, organoids were released from Cultrex by repetitive pipetting in cold PBS using a 1 ml pipette tip and transferred to a 15 ml conical tube. After centrifugation at 300× g for 5 minutes at 4°C, the supernatant was removed, and the pellet was resuspended by repeated pipetting with 1 ml Advanced DMEM/F12 (Cat. No. 12634010, Gibco). The cells were washed with an additional 1 ml of Advanced DMEM/F12 prior to resuspension in 80% Cultrex and 20% growth media.

### Generation and Propagation of Apical-Out EEO

To generate AO EEO, the growth medium was aspirated and Cultrex droplets were gently dislodged into the cell recovery solution (Cat. No. 354270, Corning and transferred to Eppendorf tubes using wide-bore 1 ml pipette tips. The tubes were rotated gently for 1 hour at 4°C. The organoids were pelleted down by centrifugation at 20× g for 3 min and washed with Advanced DMEM/F12. Finally, organoids were resuspended in an expansion medium supplemented with the ROCK inhibitor Y27632 (Cat. No. 1293823, PeproTech) and seeded as a suspension culture in ultra-low attachment plates (Cat. No. 3473, Corning). The suspension culture was gently agitated twice daily using a pipette to avoid clumping of organoids. The growth medium was replaced every other day, a process that involved transferring the organoids to a tube using wide-bore pipette tips, allowing them to settle for 5 min (or alternatively, centrifuging at 20 x g for 30 sec), aspirating the supernatant, resuspending the organoid pellet in growth medium, and seeding them in ultra-low attachment plates.

### Immunofluorescence Analysis

For immunofluorescence staining of organoids embedded in Cultrex, the expansion medium was aspirated and EEO were fixed using 2.5% paraformaldehyde (Cat. No. 15710, Electron Microscopy Sciences) for 20 min at room temperature (RT), followed by staining with hematoxylin solution (Cat. No. HHS32, Sigma Aldrich) for 10 min at RT. After washing with PBS, droplets were embedded in 2% low melting point agarose (Cat. No. 9012366, Fisher Scientific) and allowed to solidify at RT for 10 min and then at 4 °C for 1 hr. Droplets embedded in agarose were transferred from wells using a flat spatula and placed in cassettes for paraffin embedding and sectioning (5□µm). Suspension culture EEO were transferred to an Eppendorf tube and centrifuge at 20×g for 30 sec. The supernatant was aspirated and EEO were resuspended in 2.5% paraformaldehyde for 20 min, followed by staining with hematoxylin solution for 10 min at RT. After washing with PBS, organoids were transferred to plastic molds using wide bore pipet tip, excessive PBS was aspirated and organoids were embedded in 2% low melting point agarose and allowed to solidify at RT for 10 min and then at 4 °C for 1 hr. Organoids embedded in agarose were placed in cassettes for paraffin embedding and sectioning (5□µm). All sections were mounted on slides, baked at 60 °C for 30 min, deparaffinized in xylene, and rehydrated in a graded alcohol series. Deparaffinized sections were subjected to antigen retrieval by incubating sections in Tris-EDTA (Cat. No. 93684; Abcam) at 95 °C. All slides were blocked with 5% (v/v) normal goat serum (NGS) (Cat. No. 016201, Thermo Fisher Scientific) in PBS at RT for 1□h and incubated with primary antibodies (PGR - Cell signaling Technology 8757S; ESR1 - Cell signaling Technology 8644S; ZO1 - Invitrogen 33-9100, GFP - Abcam ab5450, ENPP3 - Sigma-Aldrich HPA043772, KI67 – Abcam ab15580) overnight at 4°C in 1% (v/v) NGS diluted in PBS. Immunofluorescence visualization was performed with Alexa 488 or 590 or 647-conjugated secondary antibodies (1:500 dilution; Cat. Nos. 112545143; 111585144; 111605144, Jackson ImmunoResearch). Sections were counterstained with Hoechst 33342 (2□μg/mL dilution; Cat. No. H3570; Invitrogen) before affixing coverslips with ProLong™ Diamond Antifade Mountant (Cat. No. 36961; Invitrogen). Images were taken with a Leica DM6 B upright microscope and Leica K8 camera using Leica Application Suite X (LAS X).

### Hormone Treatment

AO or AI EEO were either plated in Cultrex droplets or seeded as a suspension culture in ultra-low attachment plates. Both embedded or suspension culture organoids were grown for 4 days after passaging and then treated with either vehicle as a control (100% ethanol) or 10□nM estradiol-17β (E2; Cat. No. E1024; Sigma) for 2 days. Next, organoids were treated with either vehicle control or 10 nM E2 and 1 μM medroxyprogesterone acetate (Cat. No. PHR1589; Sigma) for 6□days with media changed every 2□days. Each treatment was performed in triplicate wells, and organoids derived from two individual donors were used. Following treatment, the organoids were harvested for immunofluorescence and scanning electron microscopy (SEM).

### Scanning Electron Microscopy

All reagents were purchased from Electron Microscopy Sciences and all specimen preparation was performed at the Electron Microscopy Core of the University of Missouri. Organoids were fixed in 2 % paraformaldehyde, 2 % glutaraldehyde in 100 mM sodium cacodylate buffer pH=7.35. Fixed organoids were rinsed with 100 mM sodium cacodylate buffer, pH 7.35 containing 130 mM sucrose. Secondary fixation was performed using 1 % osmium tetroxide (Ted Pella, Inc. Redding, California) in cacodylate buffer and incubated at 4°C for 1 hour, then rinsed with cacodylate buffer, and further with distilled water. A graded dehydration series was performed using ethanol. Specimens were dried using the Tousimis Autosamdri 815 Critical Point Dryer (Tousimis, Rockville, MD) and finally sputter coated with 10 nm of platinum using the EMS 150T-ES Sputter Coater. Images were acquired with a FEI Quanta 600F scanning electron microscope (FEI, Hillsboro, OR).

### Co-culture of Organoids with Bacteria

Escherichia coli-GFP (ATCC 25922 strain) were cultured overnight in LB broth. Bacteria were pelleted by centrifugation at 15,000× g for 10 minutes at 4°C, followed by two washes with ice-cold PBS. Finally, the bacterial cells were resuspended in 1 ml of organoid growth medium.

EEO were released from the Cultrex BME using the cell recovery solution, washed with Advanced DMEM/F12, and then resuspended in the organoid expansion medium. EEO were seeded in either ultra-low attachment plates to generate AO organoids or re-embedded in Cultrex BME to grow AI organoids. The organoids were allowed to grow for 4 days. Subsequently, EEO were mixed with bacterial cells by rotating for 1 hr in a tissue culture incubator. AI EEO were released from BME as described above prior to bacterial incubation. Following incubation, excess bacterial cells were removed by thorough washing with PBS. AI organoids were re-embedded in Cultrex BME, while AO organoids were cultured in ultra-low attachment plates. The organoids were allowed to recover for 24 hr prior to live imaging using a Leica DMi8 imaging system every 48 hrs. The growth medium was replenished every other day.

### Co-culture of Organoids with Mouse Blastocysts

Organoids were released from the Cultrex BME using the cell recovery solution, washed with Advanced DMEM, resuspended in the organoid expansion medium, and then seeded onto ultra-low attachment plates to generate AO EEO as described above. Organoids were treated with either vehicle or E2 for 2 days and E2+MPA for an additional 6 days prior to the addition of blastocyst to the culture system. Blastocysts were collected from 4-6 week old C57BL/6-Tg(UBC-GFP)30Scha/J mice (RRID: IMSR_JAX:004353). The mice were superovulated by intraperitoneal administration of 5 IU of PMSG followed by 5 IU of hCG after 46-48 h (Johnson et al., 1998). After hCG injection, the mice were placed with fertile males of the same strain for overnight mating. Mice with copulatory plugs were humanely euthanized 2.5 days later, and oviducts were collected into the M2 medium (Cat. No. MR-015-D, Millipore Sigma) for embryo recovery. The collected embryos were washed with preequilibrated KSOM media (Cat. No. MR-106-D, Millipore Sigma) and cultured until day 4.5. Any blastocysts that were still in zona pellucida were subjected to an acid tyrode solution (Cat. No. MR-004-D; Millipore Sigma) to remove the zona (Hirai et al., 2011). Blastocysts were then washed with KSOM media, and 15-20 blastocysts were co-cultured with ∼50 vehicle treated or hormone-treated AO organoids from two independent donors in at 37°C, 5% CO2, 5% O2 for 24 h.

## Acknowledgments

The authors would like to thank members of the Spencer and Kelleher laboratory for helpful discussions and DeAna Grant at the Electron Microscopy Core Facility at the University of Missouri for assistance with imaging.

## Supplementary Figures

Supplementary Figure 1. Characterization of human and mouse EEO (**A**) Immunofluorescence staining of F-Actin (grey) of AI or AO human endometrial epithelial organoids; scale bars, 20 µm). (**B**) Immunofluorescence staining of Ki67 (red) and ZO1 (green) of AI or AO human endometrial epithelial organoids; scale bars, 20 µm. (**C**) Polarity reversal is detected with SEM in suspension culture for 0, 1, 4 and 8 days; scale bars, 100 µm. Areas outlined in boxes are magnified to demonstrate the distribution of microvilli on apical surface; scale bars, 20 µm. (**D**) Bright-field images of mouse organoids grown in cultrex BME or in suspension culture for 4 days showing gross morphology; scale bars, 100 µm (**a, c**). Representative images of immunofluorescence staining of ZO1 on AI or AO mouse organoids and counterstained with Hoechst; scale bars, 20 µm (**b, d**).

Supplementary Figure 2. Apical-out organoids respond to estrogen treatment (**A**) Representative images of immunofluorescence staining of ESR1 (red) and ZO1 (green) of AI and AO treated with vehicle or E2. Quantification of ESR1-positive cells. (**B**) Representative images of immunofluorescence staining of KI67 (red) and ZO1 (green). Quantification of KI67-positive cells. Organoids were counterstained with Hoechst (blue); scale bars, 20 µm. Bars represent mean ± SD. Datapoints on the plots represent the different organoids. \*\**p* < 0.01 and \*\*\**p* < 0.001 (Student’s t test).

## References

Aplin, J. D. & Kimber, S. J. 2004. Trophoblast-uterine interactions at implantation. Reprod Biol Endocrinol, 2, 48.

Bartfeld, S., Bayram, T., Van De Wetering, M., Huch, M., Begthel, H., Kujala, P., Vries, R., Peters, P. J. & Clevers, H. 2015. In vitro expansion of human gastric epithelial stem cells and their responses to bacterial infection. Gastroenterology, 148, 126–136 e6.

Bartfeld, S. & Clevers, H. 2015. Organoids as Model for Infectious Diseases: Culture of Human and Murine Stomach Organoids and Microinjection of Helicobacter Pylori. J Vis Exp.

Benner, M., Ferwerda, G., Joosten, I. & Van Der Molen, R. G. 2018. How uterine microbiota might be responsible for a receptive, fertile endometrium. Hum Reprod Update, 24, 393–415.

Boggavarapu, N. R., Lalitkumar, S., Joshua, V., Kasvandik, S., Salumets, A., Lalitkumar, P. G. & Gemzell-Danielsson, K. 2016. Compartmentalized gene expression profiling of receptive endometrium reveals progesterone regulated ENPP3 is differentially expressed and secreted in glycosylated form. Sci Rep, 6, 33811.

Boretto, M., Cox, B., Noben, M., Hendriks, N., Fassbender, A., Roose, H., Amant, F., Timmerman, D., Tomassetti, C., Vanhie, A., Meuleman, C., Ferrante, M. & Vankelecom, H. 2017. Development of organoids from mouse and human endometrium showing endometrial epithelium physiology and long-term expandability. Development, 144, 1775–1786.

Boretto, M., Maenhoudt, N., Luo, X., Hennes, A., Boeckx, B., Bui, B., Heremans, R., Perneel, L., Kobayashi, H., Van Zundert, I., Brems, H., Cox, B., Ferrante, M., Uji, I. H., Koh, K. P., D’Hooghe, T., Vanhie, A., Vergote, I., Meuleman, C., Tomassetti, C., Lambrechts, D., Vriens, J., Timmerman, D. & Vankelecom, H. 2019. Patient-derived organoids from endometrial disease capture clinical heterogeneity and are amenable to drug screening. Nat Cell Biol, 21, 1041–1051.

Brunham, R. C., Gottlieb, S. L. & Paavonen, J. 2015. Pelvic inflammatory disease. N Engl J Med, 372, 2039–48.

Burney, R. O. & Giudice, L. C. 2012. Pathogenesis and pathophysiology of endometriosis. Fertil Steril, 98, 511–9.

Co, J. Y., Margalef-Catala, M., Li, X., Mah, A. T., Kuo, C. J., Monack, D. M. & Amieva, M. R. 2019. Controlling Epithelial Polarity: A Human Enteroid Model for Host-Pathogen Interactions. Cell Rep, 26, 2509–2520 e4.

Co, J. Y., Margalef-Catala, M., Monack, D. M. & Amieva, M. R. 2021. Controlling the polarity of human gastrointestinal organoids to investigate epithelial biology and infectious diseases. Nat Protoc, 16, 5171–5192.

Cools, P. 2017. The role of Escherichia coli in reproductive health: state of the art. Res Microbiol, 168, 892–901.

Fitzgerald, H. C., Dhakal, P., Behura, S. K., Schust, D. J. & Spencer, T. E. 2019. Self-renewing endometrial epithelial organoids of the human uterus. Proc Natl Acad Sci U S A, 116, 23132–23142.

Fitzgerald, H. C., Kelleher, A. M., Ranjit, C., Schust, D. J. & Spencer, T. E. 2023. Basolateral secretions of human endometrial epithelial organoids impact stromal cell decidualization. Mol Hum Reprod, 29.

Fitzgerald, H. C., Schust, D. J. & Spencer, T. E. 2021. In vitro models of the human endometrium: evolution and application for women’s health. Biol Reprod, 104, 282–293.

Giobbe, G. G., Bonfante, F., Jones, B. C., Gagliano, O., Luni, C., Zambaiti, E., Perin, S., Laterza, C., Busslinger, G., Stuart, H., Pagliari, M., Bortolami, A., Mazzetto, E., Manfredi, A., Colantuono, C., Di Filippo, L., Pellegata, A. F., Panzarin, V., Thapar, N., Li, V. S. W., Eaton, S., Cacchiarelli, D., Clevers, H., Elvassore, N. & De Coppi, P. 2021. SARS-CoV-2 infection and replication in human gastric organoids. Nat Commun, 12, 6610.

Gruber, T. M. & Mechsner, S. 2021. Pathogenesis of Endometriosis: The Origin of Pain and Subfertility. Cells, 10.

Hirai, H., Tani, T., Katoku-Kikyo, N., Kellner, S., Karian, P., Firpo, M. & Kikyo, N. 2011. Radical acceleration of nuclear reprogramming by chromatin remodeling with the transactivation domain of MyoD. Stem Cells, 29, 1349–61.

Hirako, I. C., Antunes, M. M., Rezende, R. M., Hojo-Souza, N. S., Figueiredo, M. M., Dias, T., Nakaya, H., Menezes, G. B. & Gazzinelli, R. T. 2022. Uptake of Plasmodium chabaudi hemozoin drives Kupffer cell death and fuels superinfections. Sci Rep, 12, 19805.

Johnson, J., Bierle, B. M., Gallicano, G. I. & Capco, D. G. 1998. Calcium/calmodulin-dependent protein kinase II and calmodulin: regulators of the meiotic spindle in mouse eggs. Dev Biol, 204, 464–77.

Kakni, P., Lopez-Iglesias, C., Truckenmuller, R., Habibovic, P. & Giselbrecht, S. 2023. PSC-derived intestinal organoids with apical-out orientation as a tool to study nutrient uptake, drug absorption and metabolism. Front Mol Biosci, 10, 1102209.

Kelleher, A. M., Demayo, F. J. & Spencer, T. E. 2019. Uterine Glands: Developmental Biology and Functional Roles in Pregnancy. Endocr Rev, 40, 1424–1445.

Lessey, B. A., Killam, A. P., Metzger, D. A., Haney, A. F., Greene, G. L. & Mccarty, K. S., JR. 1988. Immunohistochemical analysis of human uterine estrogen and progesterone receptors throughout the menstrual cycle. J Clin Endocrinol Metab, 67, 334–40.

Mitchell, C. M., Haick, A., Nkwopara, E., Garcia, R., Rendi, M., Agnew, K., Fredricks, D. N. & Eschenbach, D. 2015. Colonization of the upper genital tract by vaginal bacterial species in nonpregnant women. Am J Obstet Gynecol, 212, 611 e1–9.

Moreno, I., Codoner, F. M., Vilella, F., Valbuena, D., Martinez-Blanch, J. F., Jimenez-Almazan, J., Alonso, R., Alama, P., Remohi, J., Pellicer, A., Ramon, D. & Simon, C. 2016. Evidence that the endometrial microbiota has an effect on implantation success or failure. Am J Obstet Gynecol, 215, 684–703.

Murphy, A. R., Campo, H. & Kim, J. J. 2022. Strategies for modelling endometrial diseases. Nat Rev Endocrinol.

Murphy, C. R. 2004. Uterine receptivity and the plasma membrane transformation. Cell Res, 14, 259–67.

Nash, T. J., Morris, K. M., Mabbott, N. A. & Vervelde, L. 2021. Inside-out chicken enteroids with leukocyte component as a model to study host-pathogen interactions. Commun Biol, 4, 377.

Rawlings, T. M., Makwana, K., Taylor, D. M., Mole, M. A., Fishwick, K. J., Tryfonos, M., Odendaal, J., Hawkes, A., Zernicka-Goetz, M., Hartshorne, G. M., Brosens, J. J. & Lucas, E. S. 2021. Modelling the impact of decidual senescence on embryo implantation in human endometrial assembloids. Elife, 10.

Simintiras, C. A., Dhakal, P., Ranjit, C., Fitzgerald, H. C., Balboula, A. Z. & Spencer, T. E. 2021. Capture and metabolomic analysis of the human endometrial epithelial organoid secretome. Proc Natl Acad Sci U S A, 118.

Strauss, J. F. & Barbieri, R. L. 2019. Yen & Jaffe’s reproductive endocrinology : physiology, pathophysiology, and clinical management, Philadelphia, PA, Elsevier.

Turco, M. Y., Gardner, L., Hughes, J., Cindrova-Davies, T., Gomez, M. J., Farrell, L., Hollinshead, M., Marsh, S. G. E., Brosens, J. J., Critchley, H. O., Simons, B. D., Hemberger, M., Koo, B. K., Moffett, A. & Burton, G. J. 2017. Long-term, hormone-responsive organoid cultures of human endometrium in a chemically defined medium. Nat Cell Biol, 19, 568–577.

Wang, Y., Nicholes, K. & Shih, I. M. 2020. The Origin and Pathogenesis of Endometriosis. Annu Rev Pathol, 15, 71–95.

Wijesekara, P., Patel, K. Z., Otto, E. L., Campbell, P. G. & Ren, X. 2023. Protocol to engineer apical-out airway organoids using suspension culture of human airway basal stem cell aggregates. STAR Protoc, 4, 102154.

