## Supplemental figures 1 and 2 for "Development of Polarity-Reversed Endometrial Epithelial Organoids"

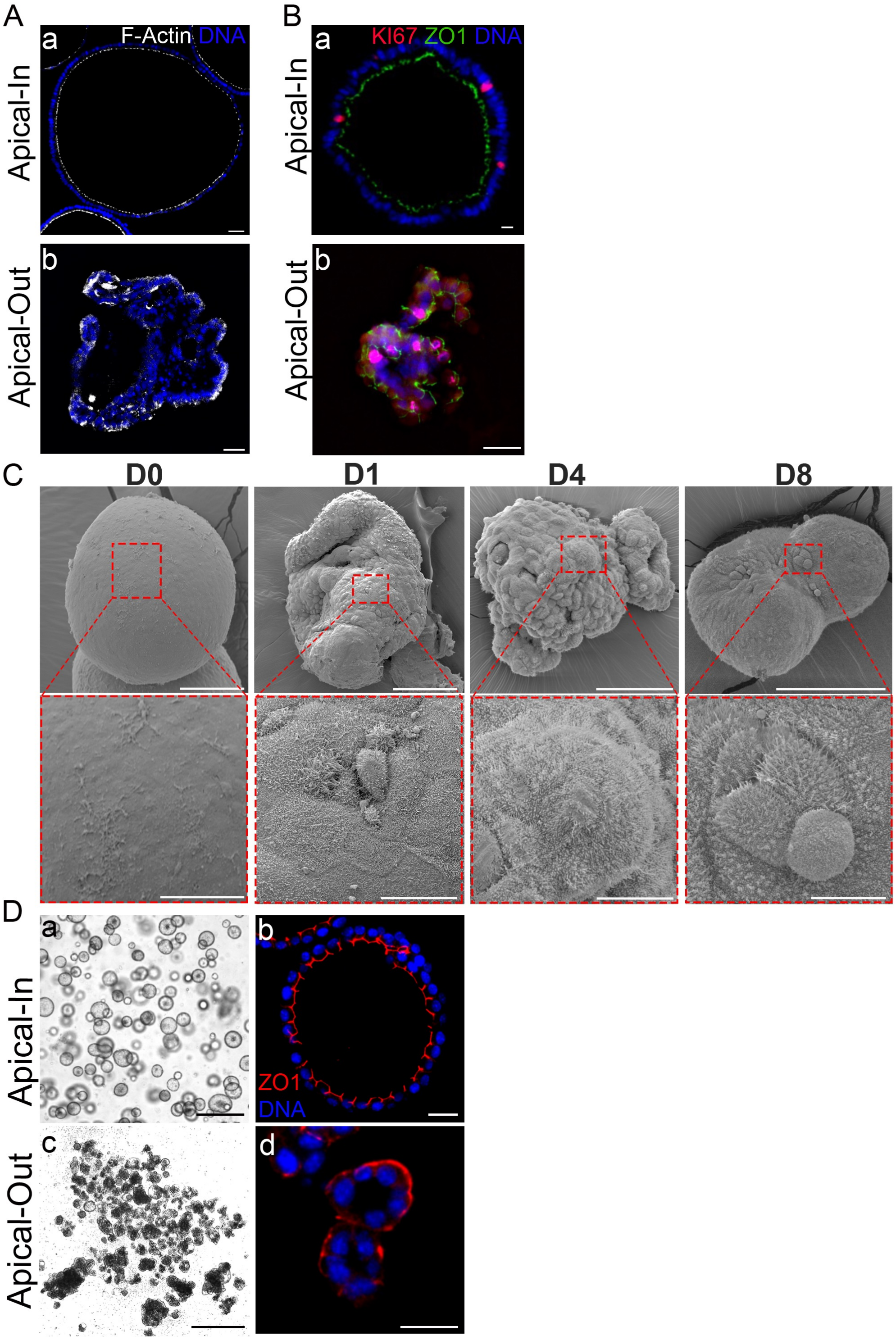


**Supplementary Figure 1. Characterization of human and mouse EEO.** (**A**) Immunofluorescence staining of F-Actin (grey) of AI or AO human endometrial epithelial organoids; scale bars, 20 μm). (**B**) Immunofluorescence staining of Ki67 (red) and ZO1 (green) of AI or AO human endometrial epithelial organoids; scale bars, 20 μm. (**C**) Polarity reversal is detected with SEM in suspension culture for 0, 1, 4 and 8 days; scale bars, 100 μm. Areas outlined in boxes are magnified to demonstrate the distribution of microvilli on apical surface; scale bars, 20 μm. (**D**) Bright-field images of mouse organoids grown in cultrex BME or in suspension culture for 4 days showing gross morphology; scale bars, 100 μm (**a**, **c**). Representative images of immunofluorescence staining of ZO1 on AI or AO mouse organoids and counterstained with Hoechst; scale bars, 20 μm (**b, d**).

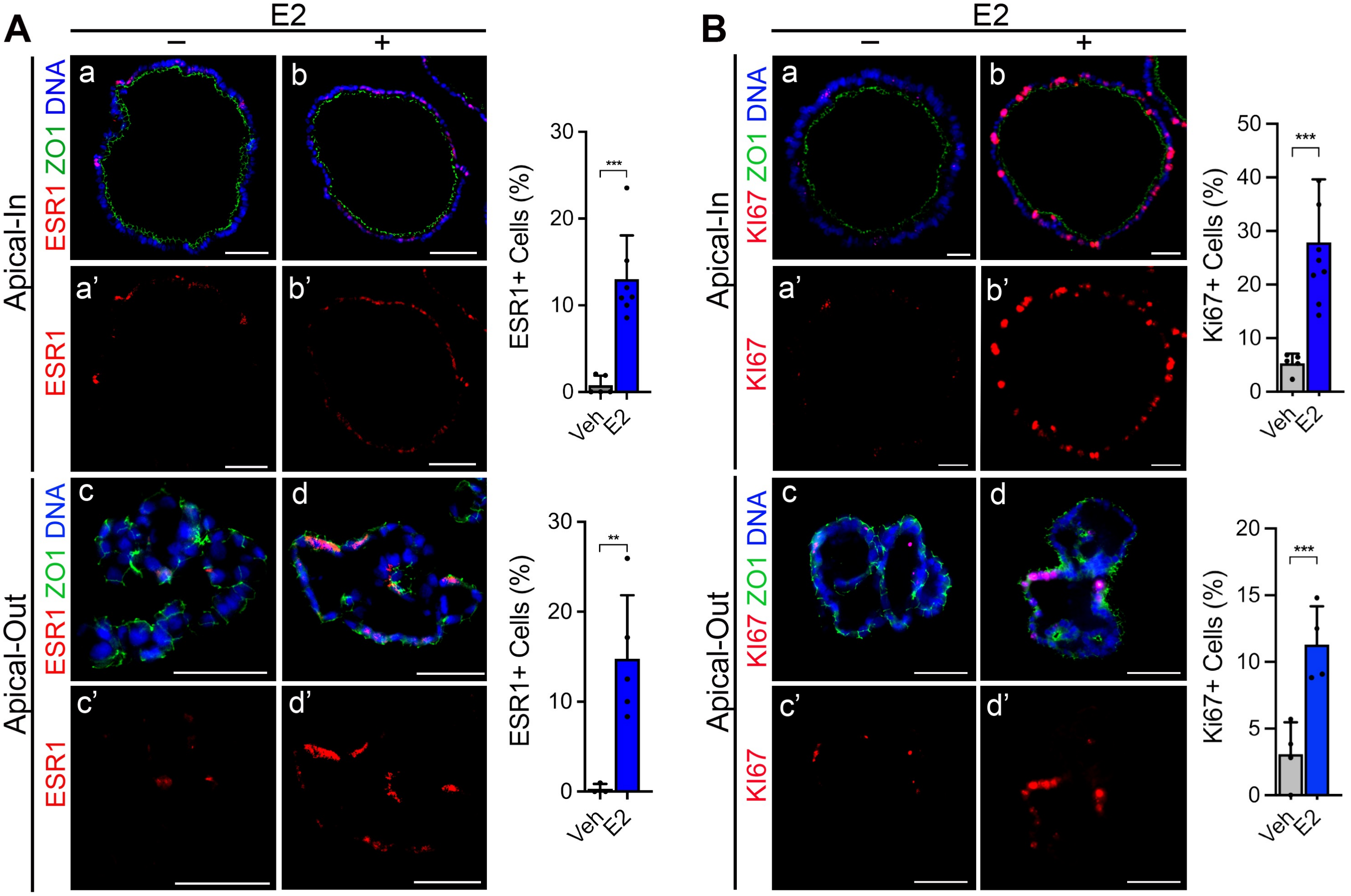


**Supplementary Figure 2. Apical-out organoids respond to estrogen treatment.** (**A**) Representative images of immunofluorescence staining of ESR1 (red) and ZO1 (green) of AI and AO treated with vehicle or E2. Quantification of ESR1-positive cells. (**B**) Representative images of immunofluorescence staining of KI67 (red) and ZO1 (green). Quantification of KI67-positive cells. Organoids were counterstained with Hoechst (blue); scale bars, 20 μm. Bars represent mean ± SD. Datapoints on the plots represent the different organoids. ***p* < 0.01 and ****p* < 0.001 (Student's t test).
